## Supplementary Figures for "Snf1/AMPK fine-tunes TORC1 signaling in response to glucose starvation"

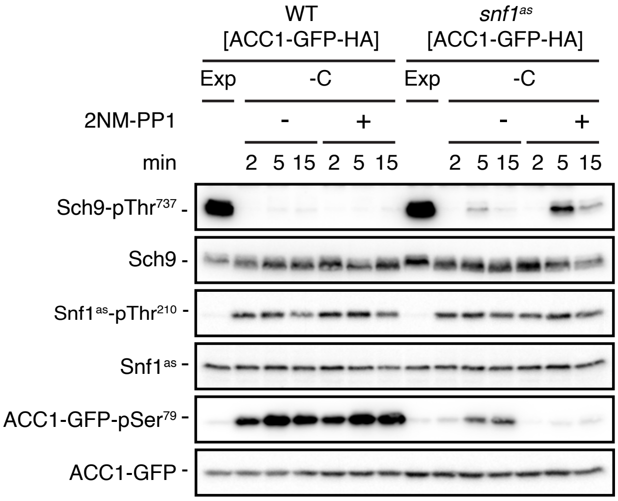


**Supplementary figure 1**

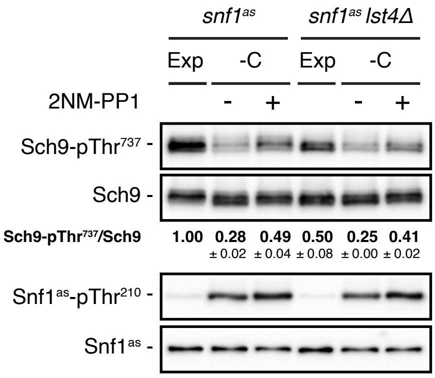


**Supplementary figure 2**

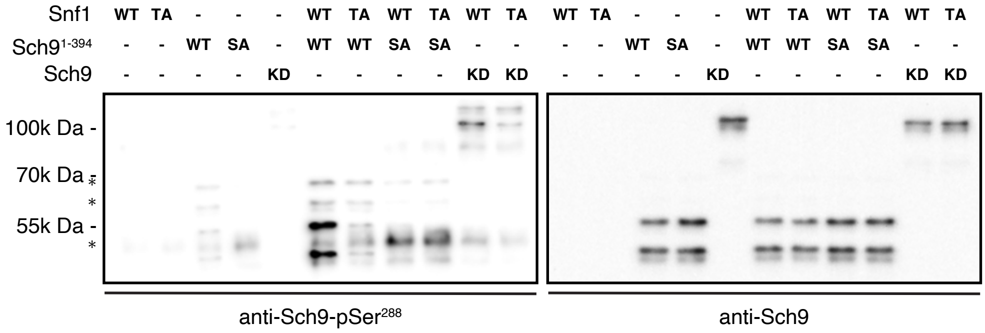


**Supplementary figure S3**

**Supplementary figure 1:** 2NM-PP1 treatment inhibits Snf1$^{as}$ activity *in vivo*. WT and ATP-analogue-sensitive *snf1$^{as}$* cells expressing a plasmid-encoded synthetic reporter of Snf1 activity that is based on a rat ACC1 peptide (ACC1-GFP; (Deroover et al., 2016)) were grown exponentially (Exp) and then starved for 2, 5, and 15 min for glucose (-C) and treated with vehicle (-; DMSO) or 2NM-PP1. Immunoblot analyses were performed as in Fig. 1A, except that anti-GFP and anti-ACC1-pSer$^{79}$ were additionally used to detect the levels of anti-ACC1-GFP and the phosphorylation state of the Snf1/AMPK target residue in ACC1-GFP that corresponds to Ser$^{79}$ in rat ACC1 (n=3).

**Supplementary figure 2:** Lst4 does not mediate TORC1 reactivation in glucose-starved, Snf1-inhibited cells. Exponentially growing *snf1$^{as}$* or *snf1$^{as}$ lst4Δ* cells (Exp) were starved for glucose (-C; 10 min) in the presence of vehicle (-; DMSO) or 2NM-PP1 (+), and analyzed as in Fig 1A. The mean TORC1 activities (*i.e.* Sch9-pThr$^{737}$/Sch9) were quantified, normalized relative to exponentially growing *snf1$^{as}$* cells (set to 1.0), and shown below the Sch9 input blot (n=3; ± SEM).

**Supplementary figure 3:** Snf1 *in vitro* kinase assays. The assays were performed as in Fig. 5C, except for the use of cold ATP and the inclusion of a full-length kinase-dead Sch9 variant (Sch9$^{KD}$: KD) as substrate. In addition, phosphorylation of Sch9-Ser$^{288}$ and the levels of Sch9$^{KD}$ or Sch9$^{1\text{-}394}$ variants were assayed by immunoblot analyses using specific anti-Sch9-pSer$^{288}$ (left blot) and anti-Sch9 (right blot) antibodies, respectively (n=2). Nomenclature: wild-type Snf1 complex: WT; kinase-inactive Snf1$^{T210A}$ complex: TA; N-terminal wild-type fragment of Sch9 (encompassing the N-terminal 394 amino acids (Sch9$^{1\text{-}394}$): WT; and Sch9$^{1\text{-}394}$ harboring the Ser$^{288}$-to-Ala mutation: SA.
