## Supplementary Tables for "Snf1/AMPK fine-tunes TORC1 signaling in response to glucose starvation"

**Table S1. Strains used in this study.**

| Strain | Genotype | Source | Figure |
|---|---|---|---|
| BY4741 | *MAT*a*; his3Δ1, leu2Δ0, met15Δ0, ura3Δ0* | Euroscarf | |
| YL515 | [BY4741] *MATα; his3Δ1, leu2Δ0, ura3Δ0* | (Binda et al., 2009) | 1A; 1E; 1F; 6C; 6E; S1 |
| MC037 | [YL515] *MATα; snf11Δ::HIS3MX6* | This study | 1A; 1F |
| MC012 | [YL515] *MATα;* $snf1^{as}$ | This study | 1A; 1C; 1F; 2A; 2C; 2E; 4F; 5D; 5F; 6A; 6D; 6E; S1; S2 |
| MC158 | [YL515] *MATα; reg1Δ::kanMX* | This study | 1E; 1F |
| MB32 | [YL515] *MATα; gtr1Δ::kanMX* | (Binda *et al.*, 2009) | 1F |
| Snf1-TAP | [BY4741] *MAT*a*; SNF1-TAP:HIS3* | Open Biosystems (Powis et al., 2015) | 4D; 5C; S3 |
| MJ5682 | [YL515] *MATα; arg4Δ::hisMX4 lys2Δ::hphNT* | (Hu et al., 2019) | |
| NIC078 | [MJ5682] *MATα;* $snf1^{as}$ | This study | 3A |
| NIC103 | [BY4741] *MAT*a*;* $snf1^{T210A}$*-TAP:HIS3* | This study | 4D; 5C; S3 |
| MC086 | [Snf1-TAP] *MAT*a*; SNF4-GFP:kanMX* | This study | 4E; 5E |
| MC013 | [MC012] *MATα; pib2Δ::HIS3MX* | This study | 4F; 6E |
| MC058 | [MC012] *MATα;* $pib2^{SASA}$ | This study | 4F; 6A; 6D; 6E |
| MC059 | [MC012] *MATα;* $pib2^{SESE}$ | This study | 4F; 6A; 6D; 6E |
| MC145 | [MC012] *MATα;* $PIB2\text{-}myc_{13}$*:kanMX* | This study | 4I |
| MC152 | [MC058] *MATα;* $pib2^{SASA}\text{-}myc_{13}$*:kanMX* | This study | 4I |
| MC153 | [MC059] *MATα;* $pib2^{SESE}\text{-}myc_{13}$*:kanMX* | This study | 4I |
| MC154 | [MC012] *MATα;* $KOG1\text{-}HA_{3}$*:HIS3MX* | This study | 4H |
| MC155 | [MC145] *MATα;* $KOG1\text{-}HA_{3}$*:HIS3MX* | This study | 4H |
| MC156 | [MC152] *MATα;* $KOG1\text{-}HA_{3}$*:HIS3MX* | This study | 4H |
| MC157 | [MC153] *MATα;* $KOG1\text{-}HA_{3}$*:HIS3MX* | This study | 4H |
| MC029 | [MC012] *MATα;* $sch9^{S288A}$ | This study | 5D; 5F; 6A; 6D; 6E |
| MC030 | [MC012] *MATα;* $sch9^{S288E}$ | This study | 5D; 5F; 6A; 6D; 6E |
| MC146 | [MC058] *MATα;* $sch9^{S288A}$ | This study | 6A; 6D; 6E |
| MC144 | [MC059] *MATα;* $sch9^{S288E}$ | This study | 6A; 6D; 6E |
| MC021 | [MC012] *MATα; lst4Δ::HIS3MX* | This study | S2 |

**Table S2. Plasmids used in this study.**

| Plasmid | Genotype | Source | Figure |
|---|---|---|---|
| pRS413 | *CEN, ARS, amp$^R$, HIS3* | (Brachmann et al., 1998) | 1A; 1C; 2A; 2C; 2E; 3A; 4F; 4H; 4I; 5D; 5F; 6A; 6C; 6D; 6E; S1; S2 |
| pRS415 | *CEN, ARS, amp$^R$, LEU2* | (Brachmann et al., 1998) | 1A; 1C; 2A; 2C; 2E; 3A; 4F; 4H; 4I; 5D; 5F; 6A; 6C; 6D; 6E; S1; S2 |
| pRS416 | *CEN, ARS, amp$^R$, URA3* | (Brachmann et al., 1998) | 1A; 1C; 2A; 2C; 2E; 3A; 4F; 4H; 4I; 5D; 5F; 6A; 6C; 6D; 6E; S1; S2 |
| pET-24d | *kan$^R$, T7p, lacO* | Novagen | |
| p3138 | [pET-24d] *His$_6$-PIB2$^{221\text{-}635}$* | This study | 4D; 4E |
| pMC030 | [pET-24d] *His$_6$-pib2$^{221\text{-}635,S268A}$* | This study | 4D |
| pMC031 | [pET-24d] *His$_6$-pib2$^{221\text{-}635,S309A}$* | This study | 4D |
| pMC032 | [pET-24d] *His$_6$-pib2$^{221\text{-}635,S268,S309A}$* | This study | 4D |
| YEplac195 | 2µ, *amp$^R$, URA3* | (Gietz and Sugino, 1988) | |
| pMC013 | [YEplac195] *GAL1p-SCH9$^{1\text{-}394}$-TAP* | This study | 5C; 5E; S3 |
| pMC016 | [YEplac195] *GAL1p-sch9$^{1\text{-}394,S288A}$-TAP* | This study | 5C; S3 |
| pMC017 | [YEplac195] *GAL1p-sch9$^{K441A}$-TAP* | This study | S3 |
| *pYX242-ACC1* | 2µ, *amp$^R$, LEU2, TPI1p-ACC1-GFP-HA* | (Deroover et al., 2016) | S1 |
| pRCC-K | 2µ, *amp$^R$, kanMX, ROX3p-CAS9, SNR52p* | (Generoso et al., 2016) | |
| pNIC012 | [pRCC-K] *SNR52p-SNF1$^{I132}$* (gRNA) | This study | 1A; 1C; 1F; 2A; 2C; 2E; 3A; 4F; 5D; 5F; 6A; 6D; 6E; S1; S2 |
| pNIC015 | [pRCC-K] *SNR52p-SNF1$^{T210}$* (gRNA) | This study | 4D; 5C; S3 |
| pMC005 | [pRCC-K] *SNR52p-SCH9$^{S288}$* (gRNA) | This study | 5D; 5F; 6A; 6D; 6E |
| pMC008 | [pRCC-K] *SNR52p-PIB2$^{S268}$* (gRNA) | This study | 4F; 6A; 6D; 6E |
| pMC009 | [pRCC-K] *SNR52p-PIB2$^{S309}$* (gRNA) | This study | 4F; 6A; 6D; 6E |

**Table S3. Oligonucleotides used in this study.**

| Name | Orientation | Sequence |
|---|---|---|
| *snf1*$^{I132}$ Proto-F | Forward | GAAATCATTATGGTTATAGAGTACGCCGTTTTAGAGCTAGAAATAGCAAGTTAAAATAAGG |
| *snf1*$^{I132}$ Proto-R | Reverse | GGCGTACTCTATAACCATAATGATTTCGATCATTTATCTTTCACTGCGGAG |
| *snf1*$^{I132G}$ Donor | Forward | TGATGTTATCAAATCCAAAGATGAAATCATTATGGTTGGAGAGTACGCCGGAAACGAATTGTTTGACTATATTGTTCAGA |
| *snf1*$^{T210}$ Proto-F | Forward | GGTAATTTCTTAAAGACTTCTTGGTTTTAGAGCTAGAAATAGCAAGTTAAAATAAGG |
| *snf1*$^{T210}$ Proto-R | Reverse | GAAGAAGTCTTTAAGAAATTACCGATCATTTATCTTTCACTGCGGAG |
| *sch9*$^{S288}$ Proto-F | Forward | GAAGATGATCTGTGTGTATAAGTTTTAGAGCTAGAAATAGCAAGTTAAAATAAGG |
| *sch9*$^{S288}$ Proto-R | Reverse | TTATACACACAGATCATCTTCGATCATTTATCTTTCACTGCGGAG |
| *sch9*$^{S288A}$ Donor | Reverse | TACTGAAGAGCAAGAGTTTAGCTGATCTAATTGGGAAGATGCTCTGTGTGTATAAAGAGGTTTTTTCTTCAAGTGCTCTT |
| *sch9*$^{S288E}$ Donor | Reverse | TACTGAAGAGCAAGAGTTTAGCTGATCTAATTGGGAAGATTCTCTGTGTGTATAAAGAGGTTTTTTCTTCAAGTGCTCTT |
| *pib2*$^{S268}$ Proto-F | Forward | GAATTCTAGCTCGATGTCCCAACTGGTTTTAGAGCTAGAAATAGCAAGTTAAAATAAGG |
| *pib2*$^{S268}$ Proto-R | Reverse | CAGTTGGGACATCGAGCTAGAATTCGATCATTTATCTTTCACTGCGGAG |
| *pib2*$^{S268A}$ Donor | Forward | GAAAATATTGTCGACAAGCTGACTACAACGAATTCTAGCGCGATGTCCCAACTGCGATTTGGCAACACGAACGTCATTAT |
| *pib2*$^{S268E}$ Donor | Forward | GAAAATATTGTCGACAAGCTGACTACAACGAATTCTAGCGAGATGTCCCAACTGCGATTTGGCAACACGAACGTCATTAT |
| *pib2*$^{S309}$ Proto-F | Forward | GAATTAAAATTCAGATTAGTGCTCGAAGCGTTTTAGAGCTAGAAATAGCAAGTTAAAATAAGG |
| *pib2*$^{S309}$ Proto-R | Reverse | GCTTCGAGCACTAATCTGAATTTTAATTCGATCATTTATCTTTCACTGCGGAG |
| *pib2*$^{S309A}$ Donor | Reverse | GATTTATGTTTATTGGAATTAAAATTCAGATTAGTGCTCTCAGCAGGCTGCGGTAAAAATTCCAGCGAGGGTTTCCTTAG |
| *pib2*$^{S309E}$ Donor | Reverse | GATTTATGTTTATTGGAATTAAAATTCAGATTAGTGCTCGAAGCAGGCTGCGGTAAAAATTCCAGCGAGGGTTTCCTTAG |
